# Spectral fingerprints distinguish buckling from angiogenesis in tortuous blood vessels

**DOI:** 10.64898/2026.06.28.735090

**Authors:** Hiroto Kawanaka, Takashi Miura

## Abstract

**Highlights:**

- Buckling and angiogenesis yield distinct vessel power spectra
- Dominant buckling wavelength vs *k*^−2^ angiogenic scaling
- Two-phase CA+EB model matches real retinal vessel morphology
- OCTA500 segments show morphology beyond single-mechanism models

Physiological blood vessels are generally straight, but tortuous curvature is observed under pathological conditions across spatial scales, from large arteries to retinal microvasculature and tumor-associated vessels. Here, we compare two theoretical mechanisms of curved vessel formation—mechanical buckling and angiogenic biased random walk—by analyzing the vessel centerline height function in Fourier space. We simplify the Chaplain–Anderson angiogenesis model and an Euler–Bernoulli buckling model with surrounding-tissue support, reproduce curvature numerically, and analyze the power spectra of the resulting patterns. Buckling yields a single characteristic peak in the power spectrum, whereas angiogenesis yields *k*^−2^ scaling in the low-frequency range. Mathematical analysis explains the selective growth of a dominant buckling wavelength in the buckling model and the origin of the scaling in the Chaplain–Anderson model. We further analyze OCTA500 segments using morphological descriptors (power spectrum, autocorrelation, and mean squared displacement); neither single-mechanism prediction alone accounts for the observed morphology, motivating a two-phase model in which angiogenic structure generation is followed by mechanical remodeling. Based on this two-phase hypothesis (a Chaplain–Anderson (CA)-like scaling background plus mechanical remodeling), we introduce quantitative indices—residual low-to-high wavenumber power ratio (rLHP) and slope reversal density (SRD)—to quantify Euler–Bernoulli–like buckling contributions in vessel segments. These results suggest that spectral fingerprints and complementary metrics may help distinguish mechanically driven tortuosity from angiogenesis-driven tortuosity in vascular images.

## 1. Introduction

### 1.1. Morphology of vascular networks

Vascular networks constitute the circulatory system and deliver oxygen and nutrients throughout the body (Hall, 2020). Because of functional optimization, normal vessels are usually straight between branch points, with only a few physiological exceptions (Han, 2012). The exceptions include uterine spiral arteries and intracranial cerebral vessels. Concerning uterine vessels, regression and shrinkage of the endometrium have been proposed to increase the coiling or buckling of the spiral arterioles (Massri et al., 2023). Cerebral vessels run mainly along the cortical surface and are reported to curve during gyrification (Kulenović and Dilberović, 2004; Padget, 1948).

Vascular formation is commonly described in terms of three major processes: vasculogenesis, angiogenesis, and remodeling (Sadler, 2023). Vasculogenesis is the earliest process in which vascular networks are newly formed, as observed in the yolk sac. Angiogenesis is the process by which new vessels sprout from existing vasculature, and most vessels are formed through this process. Remodeling is the process in which vessel diameters change under influences such as blood flow, producing hierarchical organization.

### 1.2. Theoretical models of vascular pattern

Mathematical models have been developed to describe the formation of vascular patterns. For vasculogenesis, a long-established model describes endothelial cells deforming the surrounding tissue, leading to a network with a characteristic length scale (Serini, 2003). For angiogenesis, the Chaplain-Anderson model assumes that endothelial tip cells at the leading edge of sprouts undergo a random walk while being chemotactically attracted to vascular endothelial growth factor (VEGF) produced in hypoxic regions (Anderson and Chaplain, 1998). For remodeling, classic models treat the vascular network as an electrical-circuit-like network, where segment properties evolve over time according to flow-dependent rules, producing hierarchy (Honda and Yoshizato, 1997; Pries et al., 1998).

### 1.3. Curved vessels

Under pathological conditions, curved vessels frequently appear (Han, 2012). For example, in patients with hypertension and related disorders, curvature can occur over multiple scales, from the aorta at large scales to coronary arteries at intermediate scales and retinal vessels at small scales (Ciurică et al., 2019). Buckling-driven structure formation has been proposed as one mechanism underlying such curved vessels (Han, 2012). In addition, angiogenesis near tumors can also generate vessels with abnormal tortuosity compared to normal vasculature (Nagy et al., 2009; Gaustad et al., 2021). The Chaplain-Anderson model of biased random walk is proposed to be the mechanism of this pattern formation (Anderson and Chaplain, 1998). However, tortuous morphologies alone do not uniquely identify the underlying mechanism, motivating quantitative diagnostics that distinguish mechanically driven buckling from angiogenesis-driven random-walk-like growth.

### 1.4. Research summary

In this study, we first simplified existing models and formulated two mechanisms of curved-structure formation—buckling and biased random walk—within a common framework as dynamics of the height function *h*(*x, t*). We then reproduced curvature formation numerically. We then compared the generated structures and showed that buckling yields a peak at a characteristic wavelength, whereas random walk yields a scaling with slope −2 in the power spectrum. We further explored the origin of these frequency characteristics through mathematical analysis. Finally, we analyzed frequency characteristics using a retinal vessel dataset.

## 2. Model

### 2.1. Chemotaxis: Chaplain-Anderson model

The Chaplain-Anderson model reproduces the formation of curved vascular structures observed in tumor angiogenesis through a combination of chemotaxis and random walk (Anderson and Chaplain, 1998) (Fig. 2). Tumors contain hypoxic regions that secrete vascular endothelial growth factor (VEGF) (Shweiki et al., 1992; Massri et al., 2023). Tip cells at the leading edge of vessels respond to the gradient of the VEGF concentration field *c*(**r***, t*) and undergo chemotactic movement while also experiencing random motion. Therefore, tip-cell motion is described by the following stochastic differential equation:

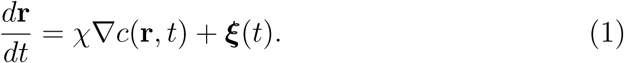

**Figure 1:**
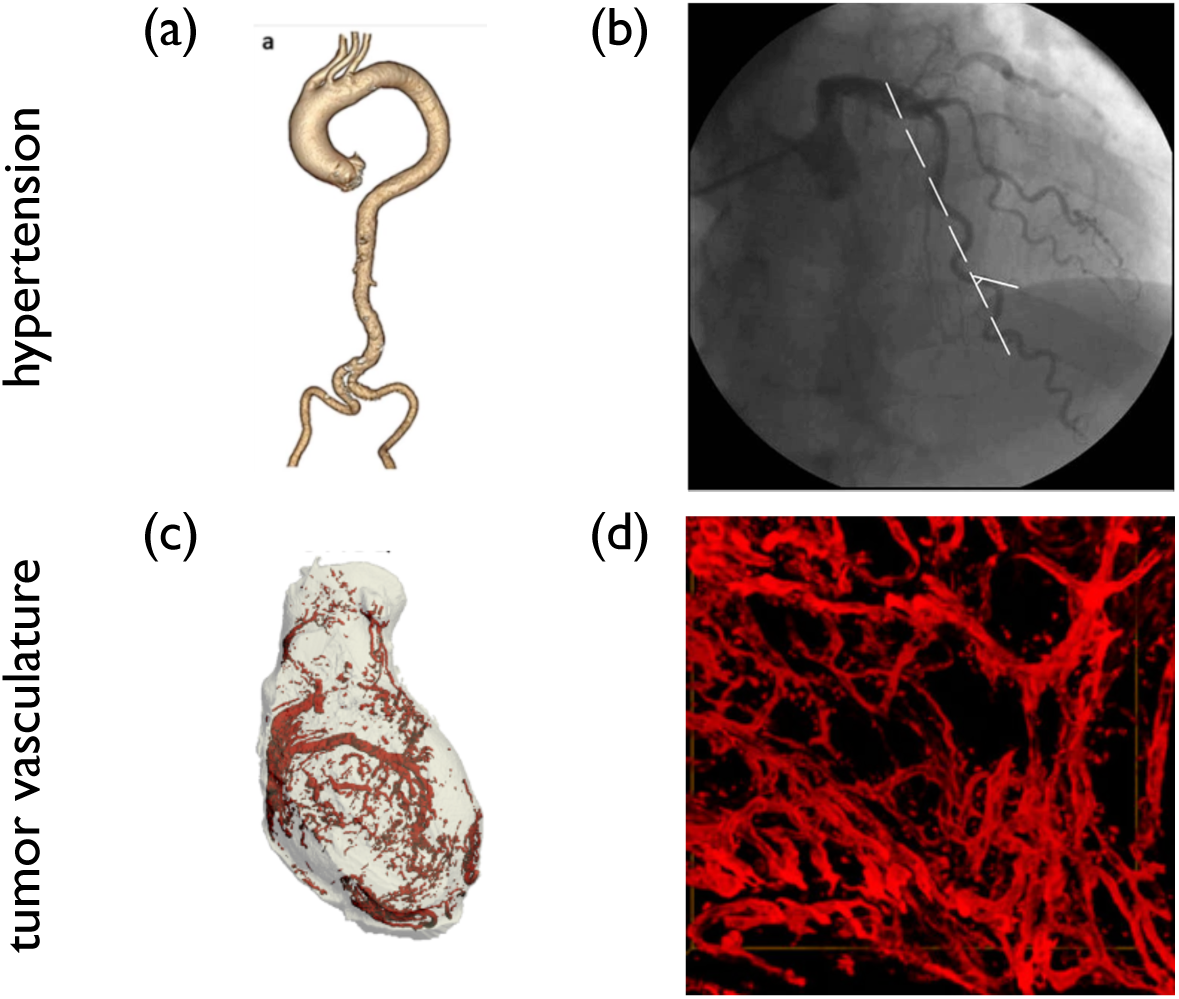
Representative examples of vascular tortuosity and abnormal vascular morphologies. (a) Aortic tortuosity associated with hypertension (adapted from (Luta et al., 2024), CC BY 4.0). (b) Coronary artery tortuosity (adapted from (Li et al., 2011), CC BY 4.0). (c,d) Tumor vasculature (adapted from (Downey et al., 2012), CC BY 4.0, and (Liu et al., 2013), CC BY 3.0). Images were cropped and relabeled for clarity and assembled as a composite figure.

**Figure 2:**
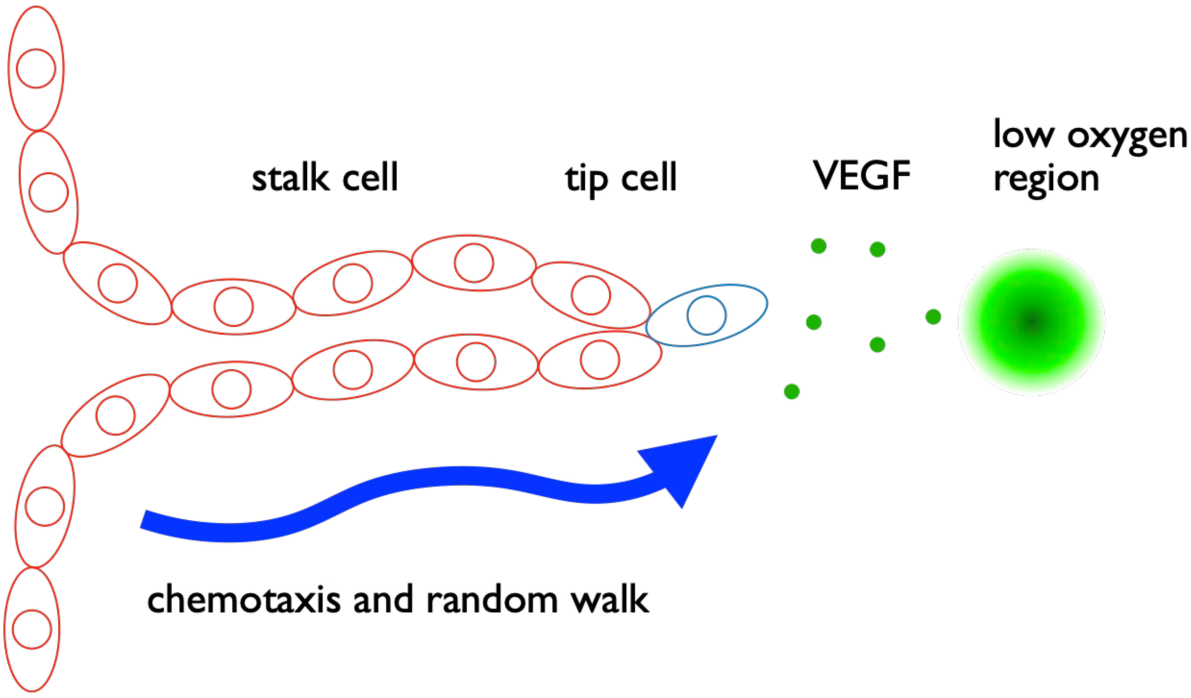
Chemotaxis-driven tumor angiogenesis (schematic based on (Anderson and Chaplain, 1998)). A migrating endothelial tip cell advances while sensing a diffusive VEGF field *c*(**r***, t*) produced by hypoxic tumor regions. Directed motion up the VEGF gradient (chemotaxis, strength *χ*) together with stochastic fluctuations can generate tortuous, irregular vessel trajectories.

Here, *χ* is the chemotactic sensitivity, and ***ξ***(*t*) is zero-mean white noise with autocorrelation

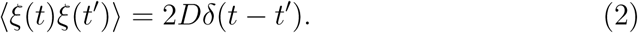

The left-hand side of the governing equation represents tip-cell velocity during angiogenesis. The first term on the right-hand side, the product of chemotactic sensitivity (*χ*) and the gradient of VEGF concentration (*c*), represents directed chemotactic migration. The second term (***ξ***) represents random tip-cell motion. The combination of these two effects induces curvature in newly formed vessels.

### 2.2. Buckling: Euler-Bernoulli model

The Euler-Bernoulli model describes how curved vessel structures emerge due to axial compressive stress applied to pre-existing vessels (Han, 2009). Axial compressive load *P* (*t*) can increase under hypertension (Han, 2012) or when age-related vessel elongation exceeds the distance between anatomical attachment sites (Han, 2012; Sugawara et al., 2008). When this compression exceeds a critical level, vessels undergo buckling. Vessels are surrounded by connective tissue, which exerts restoring forces that keep them in a straight state. In the schematic, this is represented by spring forces from the centerline (*x*-axis), although in reality, vessels receive support from connective tissue on both sides (Fig. 3). The governing equation is given by

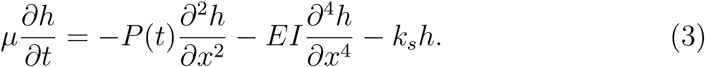

**Figure 3:**
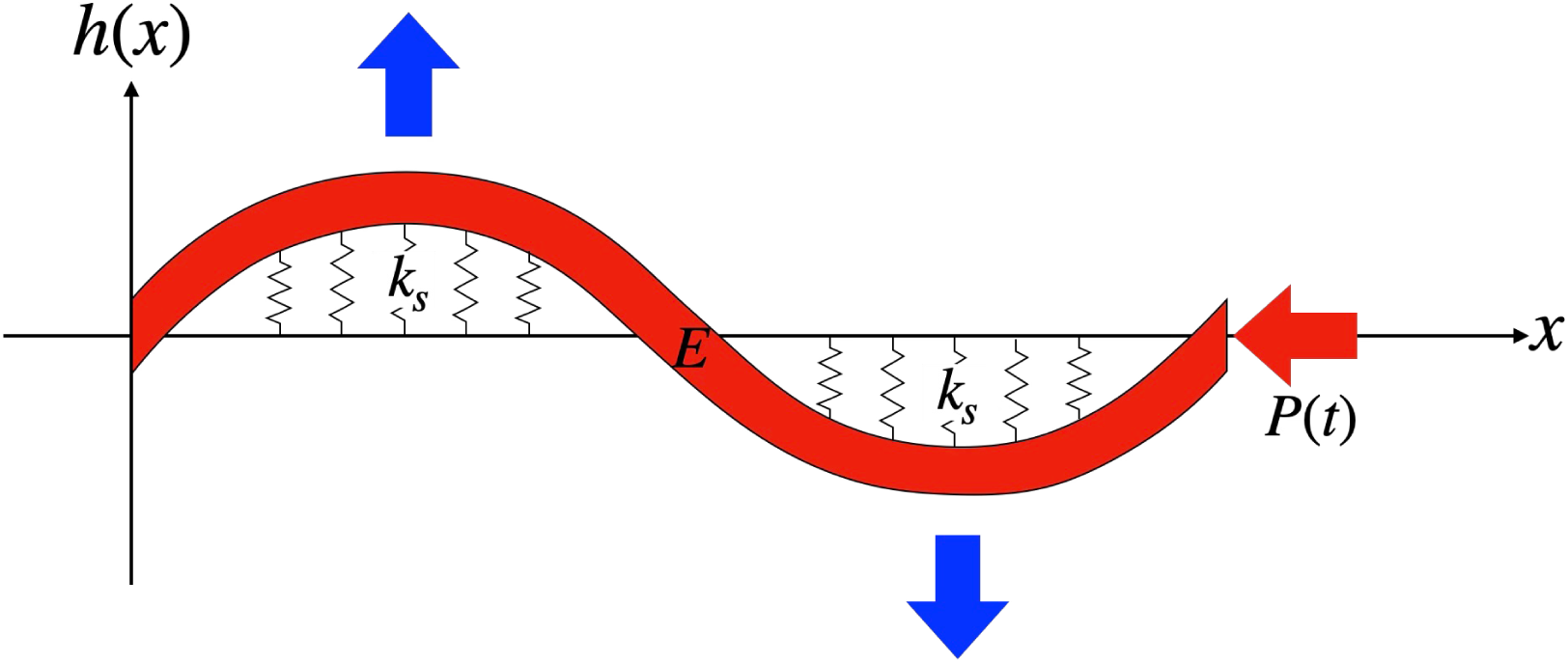
Buckling model driven by vessel elongation (schematic based on (Han, 2012)). If a vessel’s natural (stress-free) length increases due to aging, growth, or pressure-induced remodeling while its endpoints remain constrained by surrounding tissue, an effective axial compressive load *P* (*t*) builds up. Beyond a critical load, the straight configuration becomes unstable and the vessel buckles into a tortuous shape, resisted by bending stiffness *EI* and elastic support from the surrounding connective tissue (modeled by *k_s_*).

On the right-hand side, the first term is the buckling-driving force due to axial compression *P* (*t*); the second term is the elastic restoring force from vessel bending stiffness; and the third term is the elastic support from the surrounding connective tissue (Han, 2009). The left-hand side represents fric-tional resistance proportional to the transverse velocity (*µ ∂h/∂t*), balancing these elastic forces in the limit where inertial effects are negligible. When *∂h/∂t* = 0, the equation reduces to the static buckling balance considered by Han (2009). We treat *µ* as an effective friction coefficient of the vessel and surrounding tissue; its value affects the buckling onset time but not the selected wavelength in our linear stability analysis.

## 3. Materials and Methods

### 3.1. Model implementation

Two complementary mechanistic models were used to interpret vessel morphology in terms of angiogenic scaling and mechanical buckling.

#### Chemotaxis-driven angiogenesis (CA)

Capillary network morphology was represented using a Chaplain–Anderson–type angiogenesis model that generates branched vessel structures under chemotactic growth and branching rules. Synthetic CA vessel centerlines served as a reference for random-walk-like, scaling-dominated morphology. In spectral analyzes, the log–log slope of the bridge-profile power spectral density (PSD) was compared with a CA reference slope (*s*_CA_ ≈ −2.11) derived from the same chemotaxis simulation framework.

#### Euler–Bernoulli buckling (EB)

Buckling-like periodic structures were represented with the damped Euler–Bernoulli beam model described in the Model section. Final-state buckling profiles from constant and time-dependent axial-load simulations were used as reference morphologies (see Tables 1 and 2).

**Table 1:** Parameter values used in the finite-element model.

| Parameter | Symbol | Value | Description |
| --- | --- | --- | --- |
| Young's modulus of vessel | $E_{\text{vessel}}$ | $1.0 \times 10^6 \text{ N m}^{-2}$ | Material stiffness entering the bending rigidity $EI$ |
| Young's modulus of connective tissue | $E_{\text{connective}}$ | $2600 \text{ N m}^{-2}$ | Effective stiffness of the surrounding support tissue |
| Poisson's ratio | $\nu$ | 0.3 | Assumed near the commonly used range (0.3–0.4) for soft tissues |
| Mass density | $\rho$ | $1300 \text{ kg m}^{-3}$ | Assumed based on water-rich vessel-wall composition |
| Total transverse thickness | $l_x$ | 0.1 | Defines the model geometry (vessel + surrounding tissue thickness) |
| Axial vessel length | $l_y$ | 0.2 | Defines the model geometry (end-to-end distance) |
| Vessel width | $w$ | 0.01 m | Blood vessel diameter |
| Connective tissue elasticity | $k_s$ | $1.0 \times 10^{-4} \text{ N/m}^2$ | Elastic-foundation stiffness in Eq. (3) |

**Table 2:**
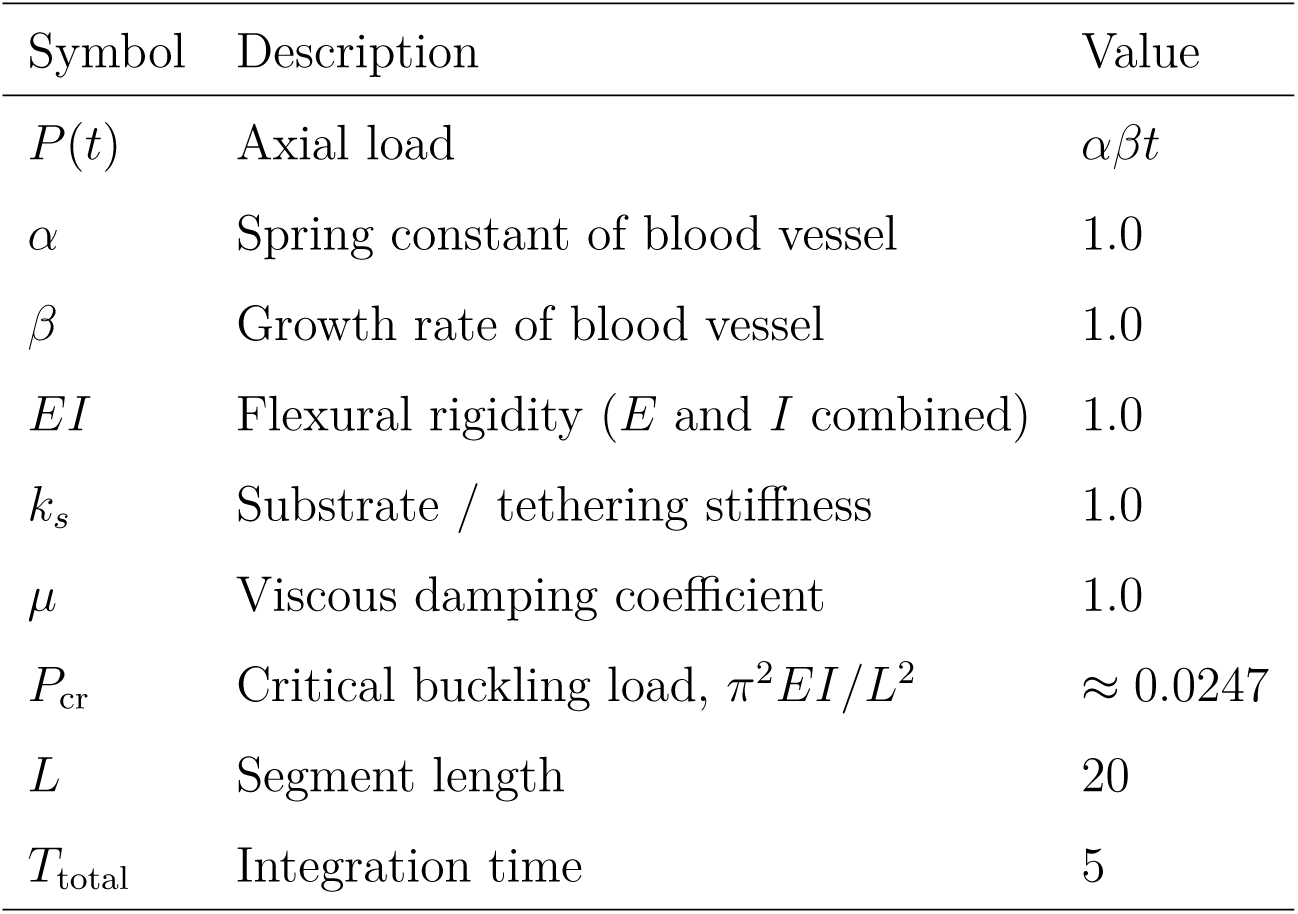
Parameters used in the Euler–Bernoulli buckling simulation with time-dependent axial load *P* (*t*) = *αβt*.

#### Synthetic reference profiles

To benchmark image-derived metrics, two reference lateral-displacement profiles were analyzed using the same pipeline as experimental segments: (i) a Brownian-bridge profile generated as a zero-mean Gaussian random walk with fixed endpoints, representing random walk-like morphology; and (ii) a final-state EB buckling profile resampled to the analysis chord length, representing buckling-dominated morphology. These references were used to interpret the image-derived rLHP and SRD metrics in terms of scaling-dominated versus buckling-dominated morphology.

### 3.2. Image analysis

#### 3.2.1. Dataset and study groups

We used the retinal vessel dataset OCTA500 (Li et al., 2024). Analyzes were performed on LargeVessel OCT angiography projection images from the GTLargeVessel category and associated disease labels, provided as grayscale bitmap files (one file per patient ID). Clinical group membership was assigned from spreadsheet labels combining disease category and age: choroidal neovascularization (CNV), retinal vein occlusion (RVO), and normal controls (age 20 years). Primary group comparisons for the rLHP/SRD analysis focused on CNV versus RVO.

All morphometric quantities were computed in pixel units, treating the image pixel as the fundamental length scale. Accordingly, spectral quantities were expressed using the corresponding pixel-based wavenumber, and comparisons emphasized dimensionless ratios and normalized metrics rather than absolute physical units; in particular, rLHP is scale-free by construction (ratio of band-limited residual power), whereas SRD is reported per pixel and is interpreted comparatively within the same imaging scale.

#### 3.2.2. Statistics

Unless stated otherwise, analyses were carried out at the segment level. Group differences were assessed with the two-sided Mann–Whitney *U* test, with the rank-biserial correlation reported as an effect size. For the primary CNV–RVO comparisons of rLHP and SRD at a single prespecified *k*_cr_, uncorrected *p*-values are reported. When multiple *k*_cr_ values were tested in the sensitivity sweep, *p*-values were additionally adjusted using the Ben-jamini–Hochberg false discovery rate procedure.

#### 3.2.3. Preprocessing and skeletonization

All image processing was implemented in Python (version 3.13.11) using OpenCV (version 4.13.0), scikit-image (version 0.26.0), and skan (version 0.13.1). Each image was read in grayscale (OpenCV imread) and binarized using Otsu’s threshold (scikit-image threshold_otsu). A 3×3 morphological closing operator with a square structuring element (OpenCV morphologyEx) was applied to suppress small gaps in the vessel mask. The binary mask was skeletonized (scikit-image skeletonize) to obtain a one-pixel-wide centerline.

#### 3.2.4. Vessel-segment extraction

Individual vessel branches were extracted as paths between junctions and endpoints using the skan library, which decomposes the skeleton graph into ordered pixel paths. The skeleton was treated as an 8-connected graph, with endpoints defined as degree-1 nodes and junctions defined as nodes with degree ≥ 3. Paths shorter than 75 pixels or longer than 150 pixels were discarded to focus on well-resolved large-vessel segments and to avoid unstable estimates from very short paths; these thresholds were chosen to be consistent with the subsequent chord-length inclusion window and fixed-length trimming. Each retained path was stored as an ordered list of pixel coordinates (*y, x*) in image space (row, column convention). For each patient image, all qualifying branch paths were treated as candidate vessel segments for downstream morphometric analysis.

Each skeleton path was converted to a chord-aligned lateral-displacement profile *h*(*x*). The path endpoints were translated to the origin and rotated so that the straight chord between endpoints aligned with the *x*-axis; the perpendicular coordinate defined *h*(*x*). Path length, chord length (start-to-end Euclidean distance), and tortuosity (path length divided by chord length) were computed from the rotated coordinates.

Segments were excluded if any of the following criteria were met:

- **Overhang:** the chord-aligned *x*-coordinate decreased along the skeleton traversal order, so that *h*(*x*) is not single-valued in *x*.
- **Chord length window:** chord length lay outside 70-110 pixels.
- **Uniform trimming failure:** the segment could not be interpolated over a fixed chord interval of length *L* = 72 pixels.

Retained segments were sorted by increasing *x*, linearly interpolated, and trimmed to *x* ∈ [*x*_0_*, x*_0_ + *L*] with *L* = 72 pixels. The bridge profile was then obtained by subtracting the linear trend from the endpoint displacements.

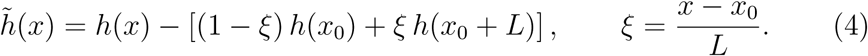

Finally, *h̃*(*x*) was resampled onto *N* = 128 uniformly spaced points in *x* for spectral and gradient-based metrics.

#### 3.2.5. Residual low-to-high wavenumber power ratio

The residual low-to-high wavenumber power ratio (rLHP) quantifies the band-limited structure in the bridge-profile spectrum after removing a scale-invariant background.

For each segment, *h̃*(*x*) was resampled onto *N* = 128 uniformly spaced points on [*x*_0_*, x*_0_ + *L*], giving a spatial step Δ*x* = *L/*(*N* − 1). The profile was mean-centered, multiplied by a Hann window *w_n_*, and transformed by a one-sided fast Fourier transform with sample spacing Δ*x*. Frequency bins *f_m_* (*m* ≥ 1) were obtained from Δ*x*, and wavenumbers were defined as *k_m_* = 2*πf_m_* (rad pixel*^−^*^1^). The periodogram was defined as the squared magnitude of each non-zero RFFT coefficient (*m* ≥ 1),

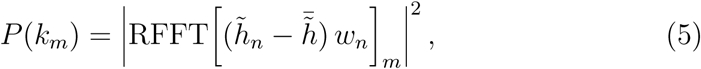

without additional normalization; the DC component (*m* = 0) was excluded.

A log–log power-law background was estimated per segment by ordinary least-squares regression over all non-zero wavenumbers *k_m_* (*m* = 1*,…,* ⌊*N/*2⌋),

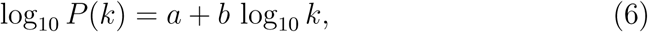

and the residual spectrum was obtained as

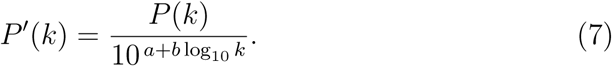

The critical wavenumber *k*_cr_ partitioned the spectrum into low- and high-wavenumber bands. The rLHP was defined as

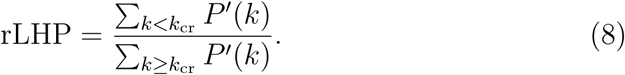

In the primary CNV–RVO comparison, *k*_cr_ = 0.5 rad pixel*^−^*^1^ was used. A sensitivity analysis was additionally performed by sweeping *k*_cr_ from 0.2 to 1.0 rad pixel*^−^*^1^ in steps of 0.1 rad pixel*^−^*^1^. Summary figures displayed rLHP on a logarithmic scale.

#### 3.2.6. Slope reversal density

Slope reversal density (SRD) quantifies fine-scale zigzag morphology in the spatial domain and is used as a complementary metric to rLHP.

On the resampled bridge profile *h̃*(*x*) at *N* = 128 points, the discrete slope was computed as Δ*h̃*/Δ*x* along uniformly spaced *x*. Adjacent slope signs were compared after assigning zero slopes the sign of the nearest non-zero neighbor. The number of sign reversals *N*_rev_ was counted along the segment. SRD was defined as the reversal count normalized by segment length,

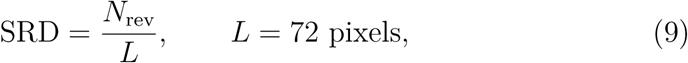

and is reported in units of pixel*^−^*^1^. In the primary analysis, no smoothing was applied before slope estimation (RAW SRD), so high-frequency fluctuations associated with random-walk-like morphology were retained. For reference, a smoothed variant was also computed in exploratory analyzes by applying a Savitzky–Golay filter (window length 15 points, polynomial order 2) to *h̃*(*x*) prior to slope estimation.

Group comparisons used the RAW SRD. A dashed reference level in summary figures indicates the SRD of a synthetic Brownian-bridge profile analyzed with the identical pipeline.

## 4. Results

### 4.1. Chemotaxis

#### 4.1.1. Numerical simulation: full model

Figure 4a shows the representative trajectories of single-vessel tip paths generated by the Chaplain-Anderson model. In this model, a vascular tip cell migrates up the VEGF gradient by chemotaxis while simultaneously receiving random fluctuations from diffusion (noise). Here, we fixed the diffusion coefficient *D* and varied the chemotactic sensitivity *χ*. When *χ* was relatively small compared with *D*, trajectories became more tortuous with larger transverse excursions; in contrast, when *χ* was large, chemotactic drift dominated, and vessels elongated more linearly along the gradient direction.

**Figure 4:**
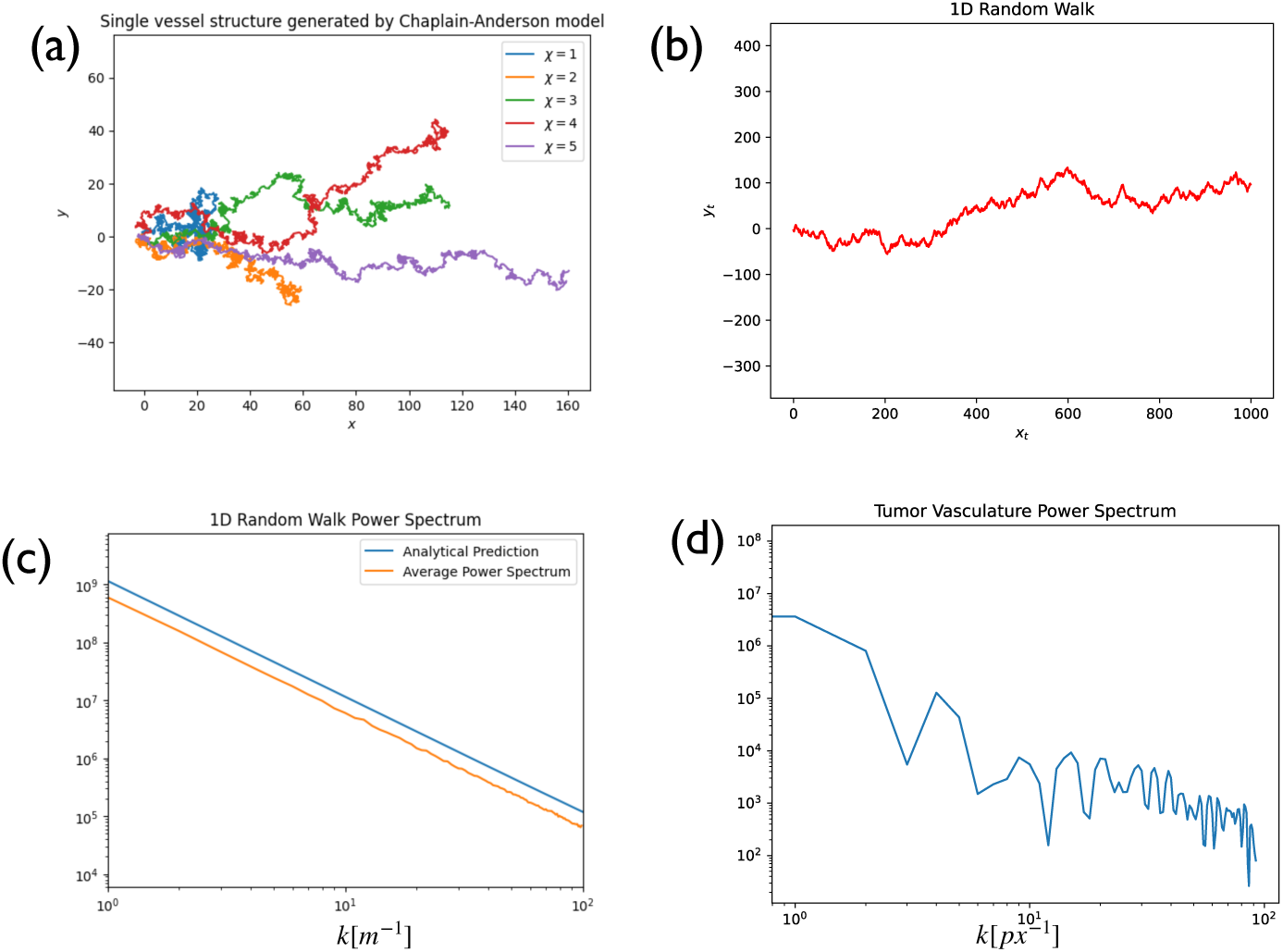
(a) Numerical results of the Chaplain-Anderson model. Vessel curvature increases when chemotactic sensitivity *χ* becomes weaker relative to diffusion coefficient *D*. (b) Numerical results of the one-dimensional random-walk model. When the random-walk component *D* is relatively small compared with chemotactic sensitivity *χ*, vessel trajectories approach a one-dimensional random walk. (c) Log-log plot comparing numerical and analytical power spectra of one-dimensional random walk. (d) Power spectrum of tumor-vessel morphology.

#### 4.1.2. Numerical simulation: one-dimensional random-walk model

Figure 4b shows numerical results for a one-dimensional random-walk approximation. The horizontal axis is the step number (time), and the vertical axis is the cumulative sum of the random increments (noise). We interpret the *x*-direction as the deterministic forward progression of the tip cell at one unit per second, i.e., drift up an approximately unidirectional VEGF gradient, while the transverse (*y*) fluctuations represent diffusion-driven randomness. This reduction corresponds to the Chaplain–Anderson regime in which chemotactic bias dominates diffusion (*χ* ≫ *D*), so that the two-dimensional biased random walk effectively projects onto a one-dimensional drift-plus-noise process (Anderson and Chaplain, 1998). Consistent with this interpretation, comparing panels (a) and (b) in Fig. 4 shows that increasing *χ* drives trajectories toward more linear, lower-tortuosity paths with smaller transverse excursions.

#### 4.1.3. Mathematical analysis

It is known that a one-dimensional random walk converges to Brownian motion under appropriate time-space rescaling (Billingsley, 1999). Brownian motion has self-affinity

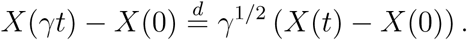

For a self-affine process *X*(*t*) with a Hurst exponent *H*, the power spectrum is

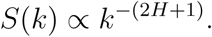

Therefore, for Brownian motion (*H* = 1*/*2), the power spectrum becomes *S*(*k*) ∝ *k*^−2^ (Falconer, 2003).

Figure 4c compares the numerically computed and analytically derived power spectra for a one-dimensional random walk. Figure 4d shows the power spectrum obtained from tumor-vessel morphology. Both exhibit *k*^−2^ scaling in the low-frequency range, indicating agreement between theory and data. These results suggest that tumor angiogenesis, governed by random-walk-like dynamics composed of both chemotactic guidance and stochastic fluctuations, shows scaling in frequency space.

### 4.2. Buckling

#### 4.2.1. Numerical simulation using finite element method

We computed buckling induced by axial compressive loading applied parallel to the axis of a straight vessel embedded in connective tissue using the finite element method implemented via NDSolve[] function in *Wolfram Language*. The vessel was surrounded by connective tissue, and a top plate covering both the vessel and tissue was placed on the right side so that the load was applied uniformly to the entire structure. Model parameters are described in Table 1.

As an initial condition, fluctuations within 1% of the vessel radius were imposed in the direction perpendicular to the vessel axis. Compressive load *P* (*t*) was determined as follows. First, the vessel ends were fixed so that the endpoint distance was constant *L*_0_(0) = *l_y_*. The natural length *L*_0_(*t*) then increased over time with a growth rate *β*[*m* · *s^−^*^1^] due to vessel-wall growth:

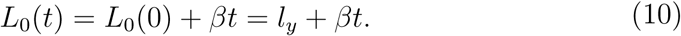

If *α*[*N* · *m^−^*^1^] denotes the spring constant of the vessel wall, the compressive stress *P* (*t*) becomes

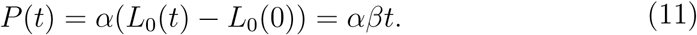

Using these parameters, the finite element simulation showed buckling and the resulting vessel curvature. For visualization, figures are displayed with a scaling factor = 1000, which exaggerates deformation relative to physical magnitude. Since this study focuses on the frequency characteristics of buckling-induced patterns and frequency is independent of this scaling factor, we adjusted the visualization to make the onset of buckling easier to identify.

We analyzed the frequency characteristics from the image data for both the numerical buckling result (Fig. 5b) and the curved coronary vessel (Fig. 1b). Log–log plots of power spectra showed a single-frequency peak in both structures (Fig. 6).

**Figure 5:**
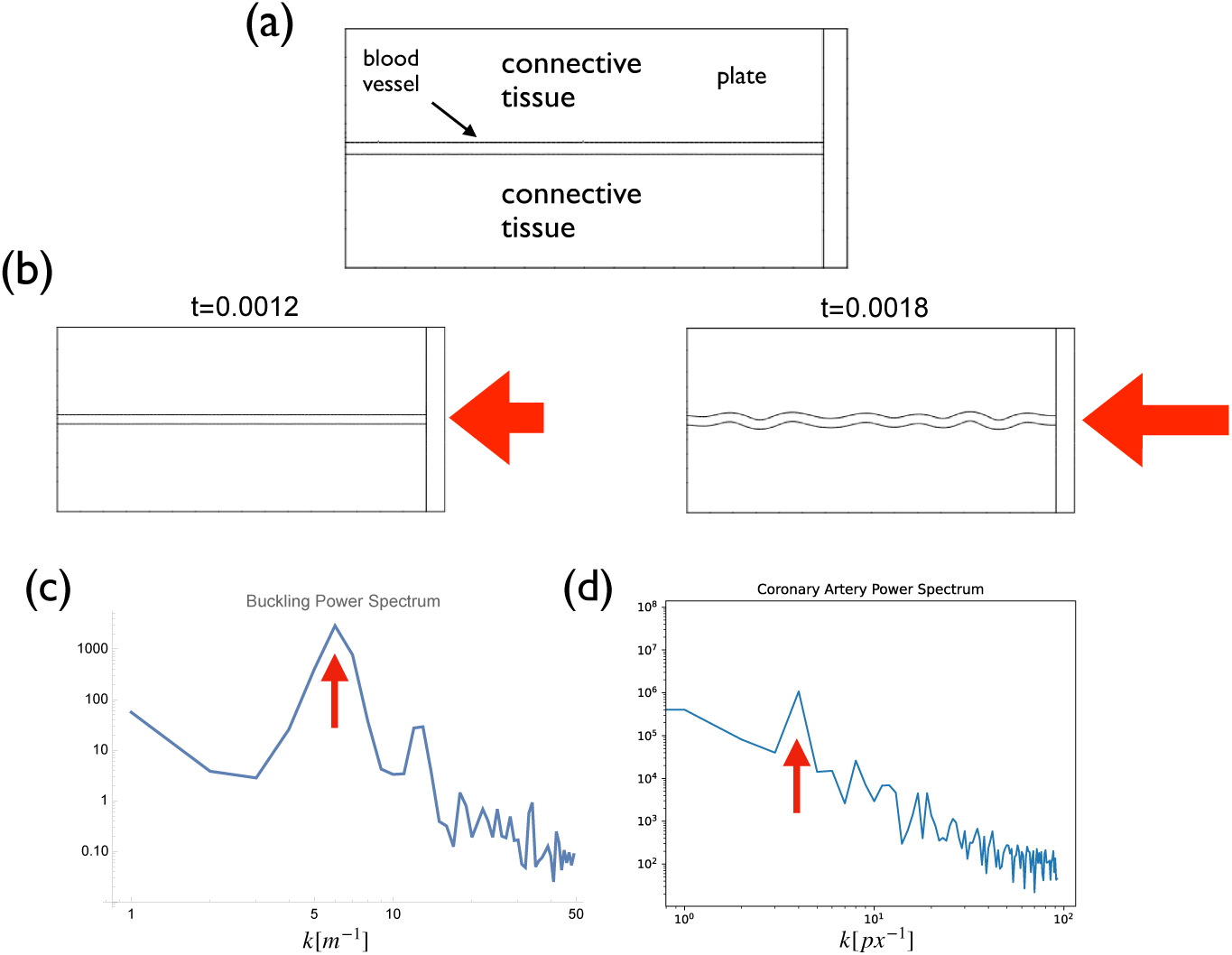
Finite-element buckling simulation. (a) The vessel was modeled as an Euler-Bernoulli-like elastic body embedded in surrounding connective tissue, represented by an elastic foundation, and compressed by a loading plate that acts on both tissue and vessel. As the effective axial load increases, the initially straight configuration becomes unstable and develops a periodic buckling mode. (b) Numerical simulation result. The computation was performed in *Wolfram Language* using NDSolve. (c-d) Log–log power spectra of (c) the finite-element buckling simulation and (d) an extracted coronary vessel centerline (Fig. 1b). In both cases, the spectrum exhibits a pronounced peak at a characteristic wavenumber (red arrows), indicating the presence of a dominant buckling wavelength rather than broadband, scale-free fluctuations.

**Figure 6:**
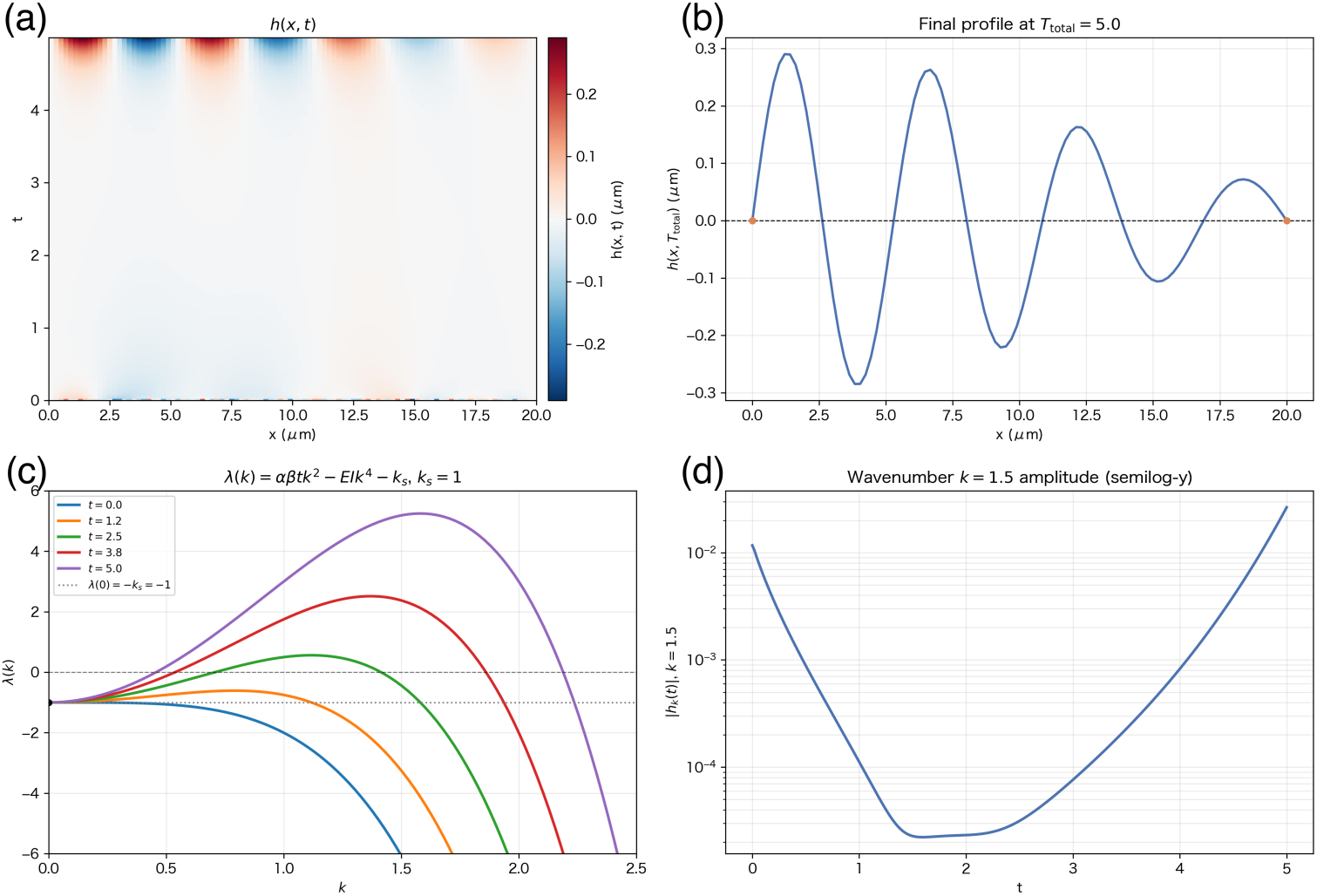
Time-dependent Euler–Bernoulli buckling under a linearly increasing axial load. Numerical solution of governing equation (3) The simulation was initialized with Gaussian noise, *h*(*x,* 0) ∼ *N*(0, 0.1^2^), and integrated to *T*_total_ = 5. **(a)** Spatiotemporal map of the transverse displacement *h*(*x, t*). **(b)** Transverse profile at the final time, *h*(*x, T*_total_). **(c)** Linear growth rate *λ*(*k*) = *αβt k*^2^ − *EI k*^4^ − *k_s_* evaluated at *t* = 0, 1.25, 2.5, 3.75, and 5; the horizontal dashed and dotted lines indicate *λ* = 0 and *λ*(0) = −*k_s_*, respectively. **(d)** Time evolution of the Fourier amplitude |*h_k_*(*t*)| at wavenumber *k* = 1.5 (semilog scale). Simulation parameters are summarized in Table 2.

#### 4.2.2. Numerical simulation using Euler-Bernoulli equation

We also used an analytically manageable governing equation (3). The parameter sets used in the model are described in Table 2. The numerical simulations reproduce the formation of a winding structure (Fig. 6).

##### Linear stability analysis

For linear stability analysis, we assume a displacement of wavenumber *k* as follows:

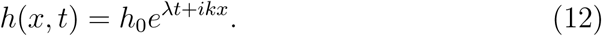

Then derivatives with respect to *x* are

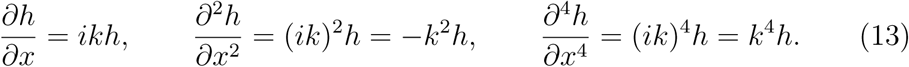

Substituting into the governing equation (3) gives the growth rate (dispersion relation):

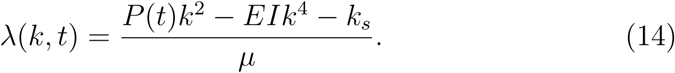

As the compressive load *P* (*t*) increases over time, the *λ*(*k*) curve shifts upward, and at a certain point, the growth rate of specific wavenumber components becomes positive at *k* = *k*_max_. This fastest-growing mode h determines the dominant observed buckling wavelength, 2*π/k*_max_.

Differentiating *λ* with respect to the wavenumber *k* gives

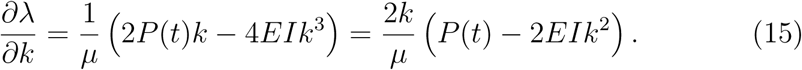

Setting this to 0, i.e.,

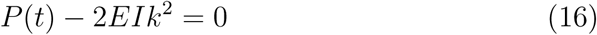

maximizes *λ*. Therefore,

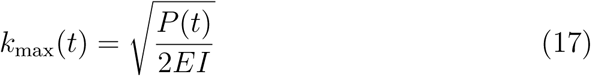

is the fastest-growing wavenumber, and the corresponding buckling wavelength is

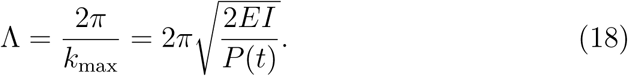

Assuming the compressive load increases linearly over time,

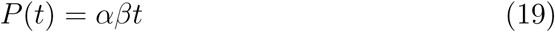

(where *α* is the spring constant and *β* is the growth rate),

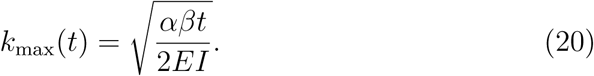

The condition for a positive growth rate (i.e., buckling onset) is *λ*(*k*_max_*, t*) *>*

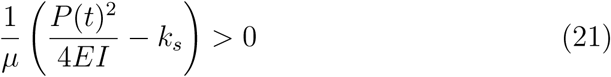

Therefore, the onset time of buckling *t_c_* is

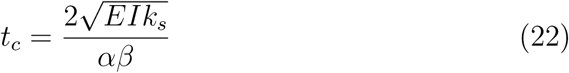

given by the above expression. The fastest growing wavenumber at *t_c_* is

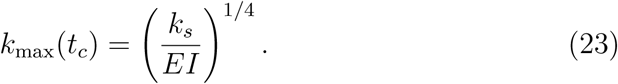

and the corresponding wavelength is

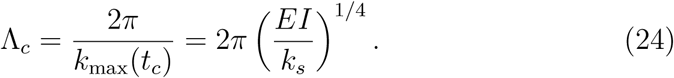

Parameters used in the numerical simulation are described in Table 2. This analysis quantitatively explains how vessels form curved structures at specific wavelength scales during temporal evolution under growth and compression. The corresponding time-evolving dispersion relation is shown in Fig. 6c.

### 4.3. Quantification of retinal vessel by power spectrum profile

#### 4.3.1. Dataset

We tested how well patterns generated by these models represent real vessel morphology using the retinal vessel dataset OCTA500 (Li et al., 2024).

Within the dataset, we used images in the GT large vessels category and grouped segments for illustration into three clinical categories: normal controls (age 20 years), choroidal neovascularization (CNV), and retinal vein occlusion (RVO) (Figure 7a). These disease groups were expected to exhibit curvature patterns associated with buckling and angiogenesis, respectively.

**Figure 7:**
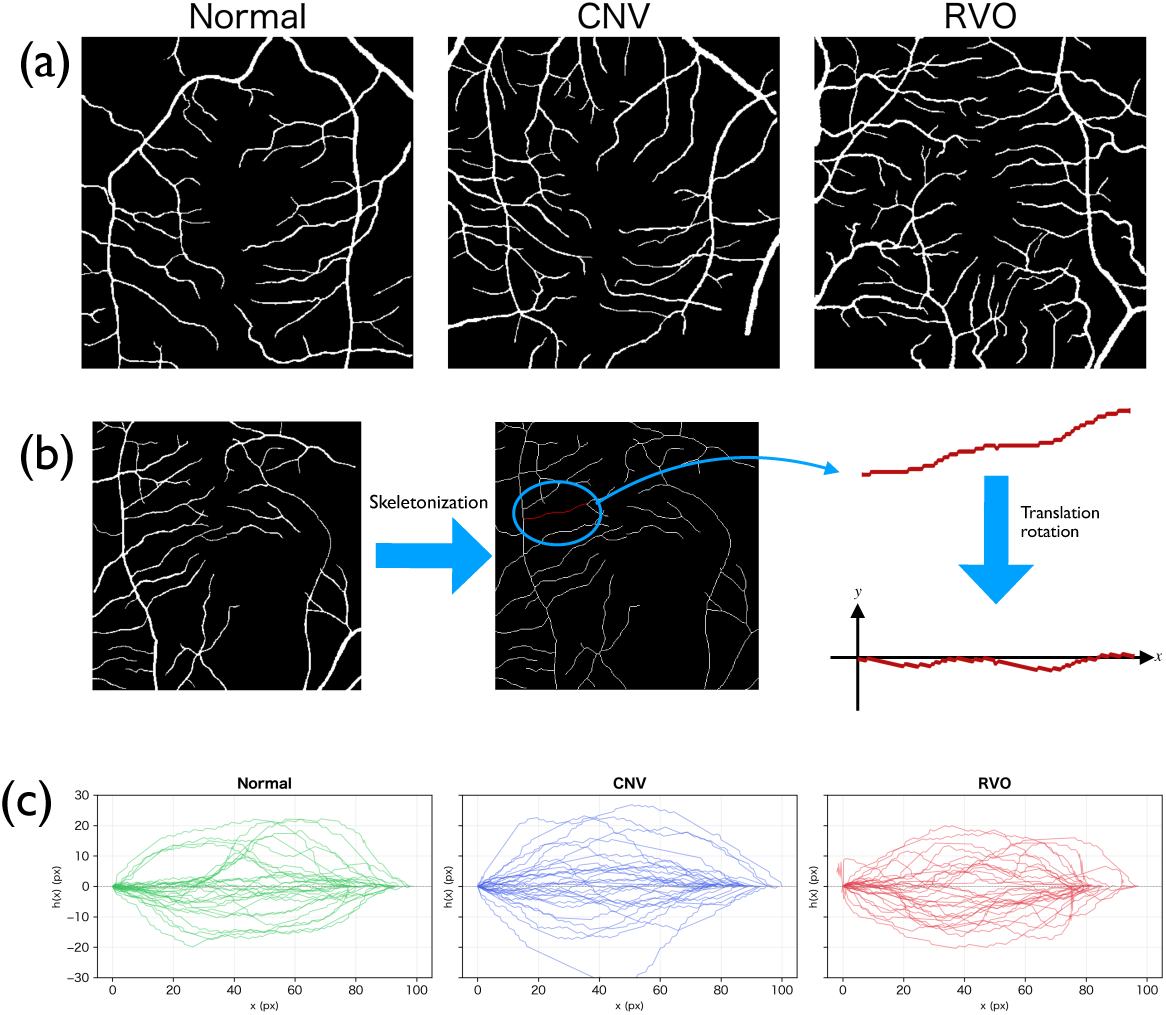
Retina vasculature images and image processing procedures. (a) Representative OCTA images of Normal, CNV, and RVO, taken from OCTA500 (Li et al., 2024). (b) schematic overview of skeletonization and chord alignment. (c) Representative aligned segment ensembles (*n* = 30 per group) for Normal, CNV, and RVO.

Retinal vein occlusion can elevate venous pressure and induce vessel dilation and elongation upstream of the obstruction; these mechanical changes may promote buckling-like tortuosity (Han, 2009). By contrast, choroidal neovascularization involves pathological neovessel growth, and we therefore treat CNV segments as a setting in which angiogenesis-driven, random-walk-like irregularity is expected to be relatively stronger than in RVO.

For each patient image, qualifying skeleton paths were converted to chord-aligned profiles *h*(*x*) using the criteria described in Methods: overhang exclusion, chord length 70–110 pixels, and successful uniform trimming to *L* = 72 pixels. Endpoints were translated and rotated so that the chord aligned with the *x*-axis, and endpoint displacements were pinned to zero after bridge detrending (Figure 7b).

#### 4.3.2. Power spectrum

First, we measured power spectra, whose theoretical characteristics are predicted by the CA and EB models described above. We expected scaling behavior if angiogenesis is the primary driver of curvature and a single peak if buckling is dominant. However, the power spectrum showed a characteristic shape with changes in curvature at two locations (Figure 9a). This feature could not be reproduced by either the CA or EB model alone.

#### 4.3.3. Autocorrelation function

Fourier transforms are often not sufficiently sensitive for detecting periodic structures in nature (Miura et al., 2000). Therefore, we assessed periodicity using autocorrelation functions. In an autocorrelation function, the value is maximal at zero lag and minimal when the phase is shifted by half a period.

With this method, a minimum was observed near lag 35 pixels in real data. Because the dataset aligns starting and ending y-coordinates, the structure can be interpreted as containing an overall half-period component, which likely explains this effect. CA showed a similar trend, whereas EB exhibited stronger periodicity, yielding multiple minima at much shorter periods.

#### 4.3.4. MSD of Brownian bridge

Next, we measured the mean square displacement (MSD, *h*(*x, t*)^2^) at each point of the curved structures. A random walk with matched start and end y-coordinates is called a Brownian bridge (Durrett et al., 1977). The theoretical mean of this system should follow a quadratic curve.

In measured MSD, the profile did not follow a symmetric quadratic curve but instead appeared slightly skewed to the right. Such a distribution cannot be reproduced by the CA or EB model alone (Figure 8).

**Figure 8:**
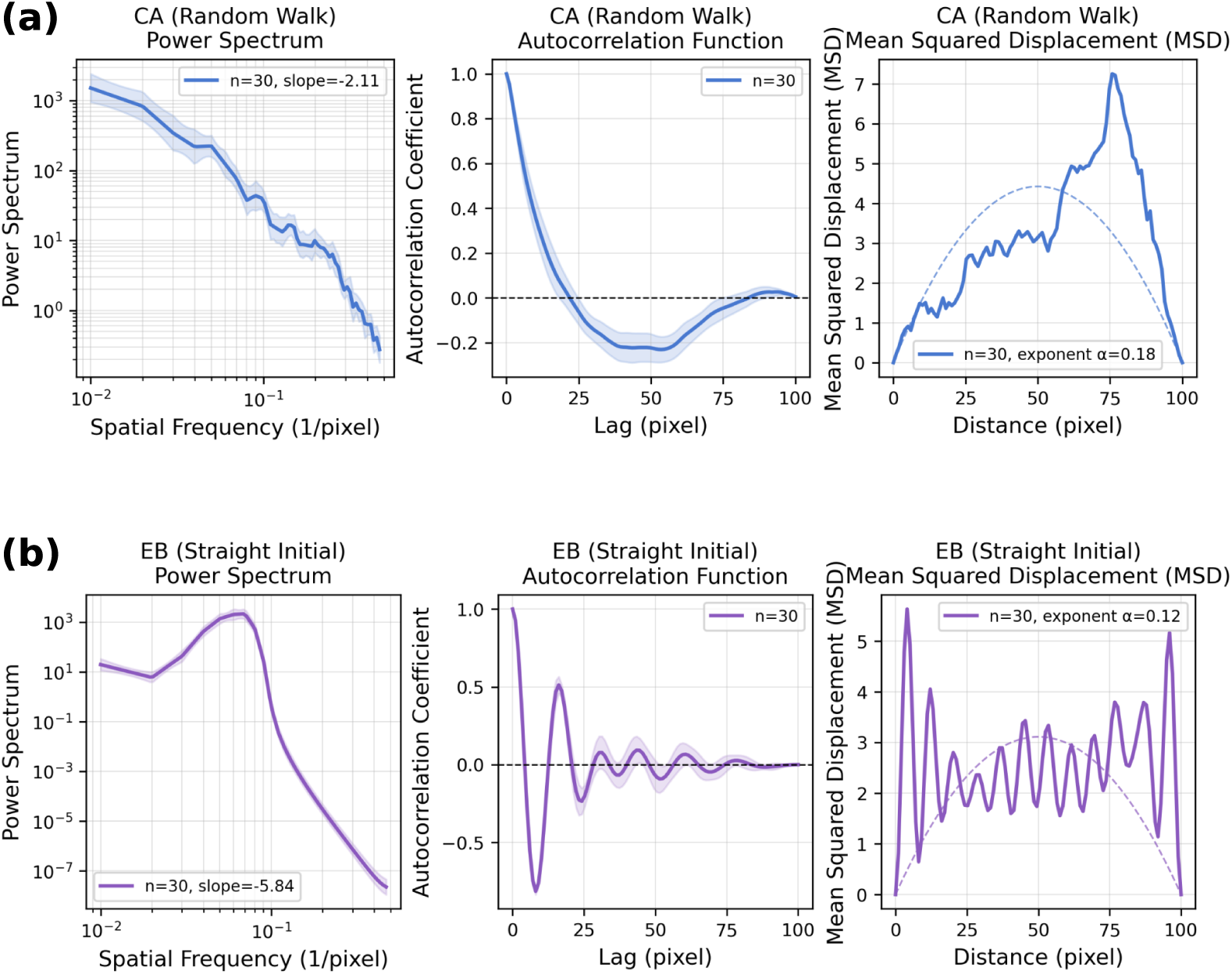
Statistical shape analysis of vessel patterns generated by CA and EB models. (a) Analysis of vessel patterns generated by the Chaplain-Anderson (CA) model. From left to right: power spectrum, autocorrelation function (ACF), and mean squared displacement (MSD). Curves show averages over *n* = 30 samples, and shaded regions indicate variability. Dashed lines are theoretical fits for MSD, with spectral slopes and MSD exponents indicated in the figure. (b) Analysis of synthetic patterns generated by the Euler-Bernoulli (EB) model, shown in the same format as A.

#### 4.3.5. Discrepancies with numerical model predictions

Comparison with observed retinal vessel morphology highlights two discrepancies: (i) the Fourier spectrum lacks a distinct dominant peak, and (ii) the log–log scaling is not well described by a single −2 slope. The missing peak likely reflects a mixture of characteristic wavelengths across segments, arising from heterogeneity in vessel diameter and local tissue mechanics. The departure from −2 scaling is consistent with the Brownian-bridge constraint: because the VEGF-gradient direction cannot be inferred reliably from morphology alone, enforcing *h*(*L*) = 0 imposes an endpoint condition that biases the statistics toward random-walk-like behavior.

### 4.4. Two-phase model

#### 4.4.1. Model definition

The Chaplain–Anderson (CA) model describes how vessel networks are generated under chemotactic growth, whereas the Euler–Bernoulli (EB) model describes how an already formed vessel can deform under axial compression and substrate tethering. Because the CA mechanism also works during the initial vessel development process, we combined them into a two-phase model: vessel structure is first generated by CA-like chemotactic growth, and the resulting centerline is subsequently relaxed and reshaped by damped EB dynamics.

*Phase 1: CA generation of the reference centerline.* In the first phase, a lateral-displacement profile *h*_0_(*x*) is generated by a simplified chemotaxis-driven random walk, which represents the initial vessel development process. The centerline advances along the chord direction with constant speed while accumulating transverse fluctuations:

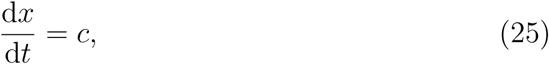

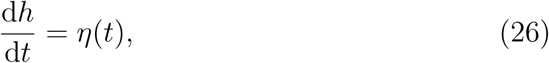

where *c >* 0 is the chemotactic drift speed and *η*(*t*) is zero-mean Gaussian white noise with variance *σ*^2^. Integration is stopped when the chord coordinate reaches a prescribed segment length *L*. The resulting time series *h_t_* is reparameterized by arc position to obtain the CA reference profile

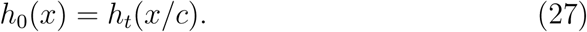

In the numerical implementation, we generated an ensemble of CA centerlines on a fixed chord length *L* = 100 using *c* = 1 and *σ_η_*= 0.5.

*Phase 2: EB remodeling on the CA-generated shape..* In the second phase, the CA profile *h*_0_(*x*) replaces the straight reference state used in the standalone EB model. Let *H*(*x, t*) denote the additional transverse displacement relative to *h*_0_(*x*), so that the total profile is *h*(*x, t*) = *H*(*x, t*) + *h*_0_(*x*). The profile evolves according to the damped EB equation

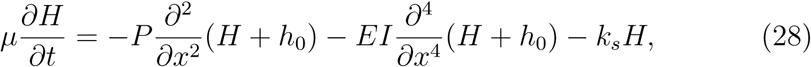

where *µ* is the viscous damping coefficient, *P* is the axial load, *EI* is flexural rigidity, and *k_s_*is substrate stiffness. In Eq. (28), axial compression and bending act on the total profile *h* = *H* + *h*_0_, whereas the elastic-foundation term −*k_s_H* damps only the mechanically induced displacement relative to the CA reference. We use the same friction coefficient *µ* as in Eq. (3). Pinned boundary conditions *H*(0*, t*) = *H*(*L, t*) = 0 were imposed so that the segment endpoints remain fixed on the CA baseline.

Before integration, *h*_0_(*x*) was interpolated onto a uniform grid of *N* = 100 points on [0*, L*] and detrended by subtracting the endpoint chord so that *h*_0_(0) = *h*_0_(*L*) = 0. The EB displacement was initialized as *H*(*x,* 0) = 0.

#### 4.4.2. Numerical simulation

Numerical simulations were performed on CA-generated centerlines taken from a precomputed ensemble of 100 vessels. For each run, one CA profile was resampled onto a uniform mesh with *L* = 100, *N* = 100, and Δ*x* = *L/*(*N* −1). Equation (28) was integrated explicitly in time with a step size Δ*t* = 0.001 to *T*_total_ = 200. Spatial derivatives were evaluated using second- and fourth-order finite differences on a padded grid with odd reflection at the boundaries. The axial load was held constant at *P* = 0.03, well below the Euler buckling load *P*_cr_ = *π*^2^*EI/L*^2^ ≈ 0.0987 for *EI* = 1. Substrate stiffness was set to *k_s_* = 10*^−^*^4^. Thirty CA vessels were analyzed per condition (random seed 42). After integration, each final profile *h*(*x, T*_total_) was converted to a bridge profile by removing the endpoint-to-endpoint linear trend. The same post-processing pipeline as for the experimental segments was applied: power spectral density (PSD), autocorrelation function (ACF), and mean squared displacement (MSD) with Brownian-bridge detrending.

Compared with the CA or EB model alone, the two-phase profiles showed better qualitative agreement with real vessel morphology. In the PSD, the model reproduced a piecewise log–log structure: a shallower slope at short wavelengths, a steeper slope at intermediate wavelengths, and a shallower slope again at long wavelengths. In the ACF, the model reproduced a profile that decreases initially and then rises toward positive correlation at larger lags. In the MSD, the model reproduced the right-skewed distribution characteristic of Brownian-bridge-like endpoint-constrained fluctuations.

### 4.5. Quantitative metrics for distinguishing the EB and CA models

Because spectral patterns are not always straightforward to classify automatically (e.g., whether a buckling-like component is present), we introduce two complementary metrics designed to quantify the contribution of the Eu-ler–Bernoulli (EB) component.

*Residual low-to-high wavenumber power ratio (rLHP).* This metric captures a localized excess (spike) in the power spectrum associated with the most unstable buckling mode. To reduce sensitivity to variability in the characteristic wavenumber (e.g., due to vessel diameter or local tissue mechanics), rLHP compares the residual power below and above a threshold wavenumber *k_c_*.

*Slope reversal density (SRD).* SRD is defined as the number of sign changes in the local slope per unit arc length. CA-dominated trajectories (random-walk-like) are expected to exhibit frequent reversals, yielding a higher SRD, whereas EB-driven buckling introduces more coherent curvature and, therefore, a lower SRD.

We applied these metrics to compare RVO and CNV segments. Based on their putative mechanisms, we expected RVO to show higher rLHP and lower SRD than CNV; the measured data followed this trend, supporting the utility of the proposed metrics (Fig. 10).

**Figure 9:**
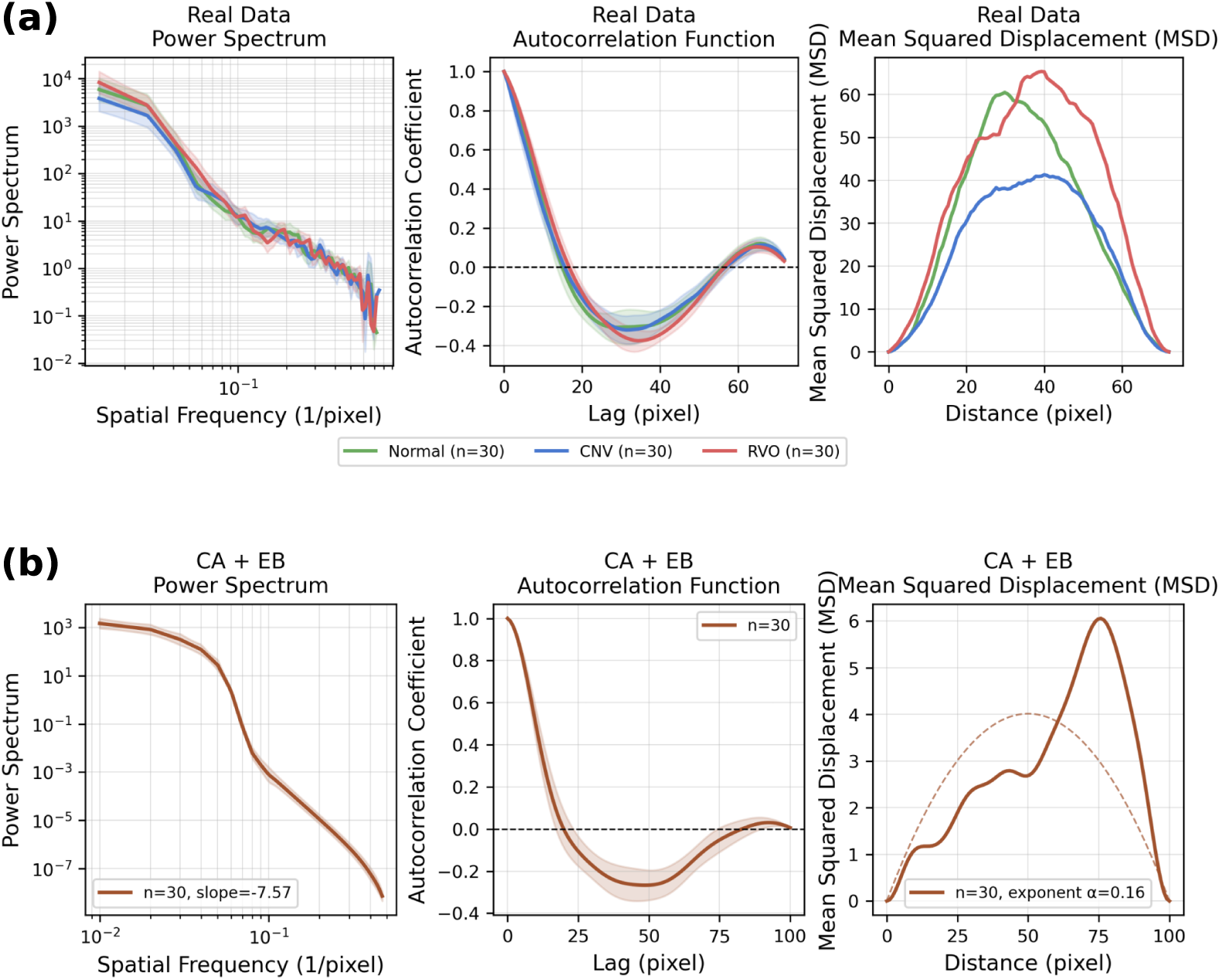
Comparison of shape analyses between real data and CA+EB synthetic vessels. (a) Analysis of segments extracted from actual retinal vessel images. From left to right: power spectrum, autocorrelation function (ACF), and mean squared displacement (MSD). For each group (Normal, CNV, RVO), curves show averages over *n* = 30 samples and shaded regions indicate variability. (b) Analysis of vessel patterns obtained by applying the Euler-Bernoulli (EB) equation to initial shapes generated by the Chaplain-Anderson (CA) model. From left to right: power spectrum, autocorrelation function (ACF), and mean squared displacement (MSD). Curves are *n* = 30 averages with variability shown by shading; dashed lines indicate theoretical MSD fits.

**Figure 10:**
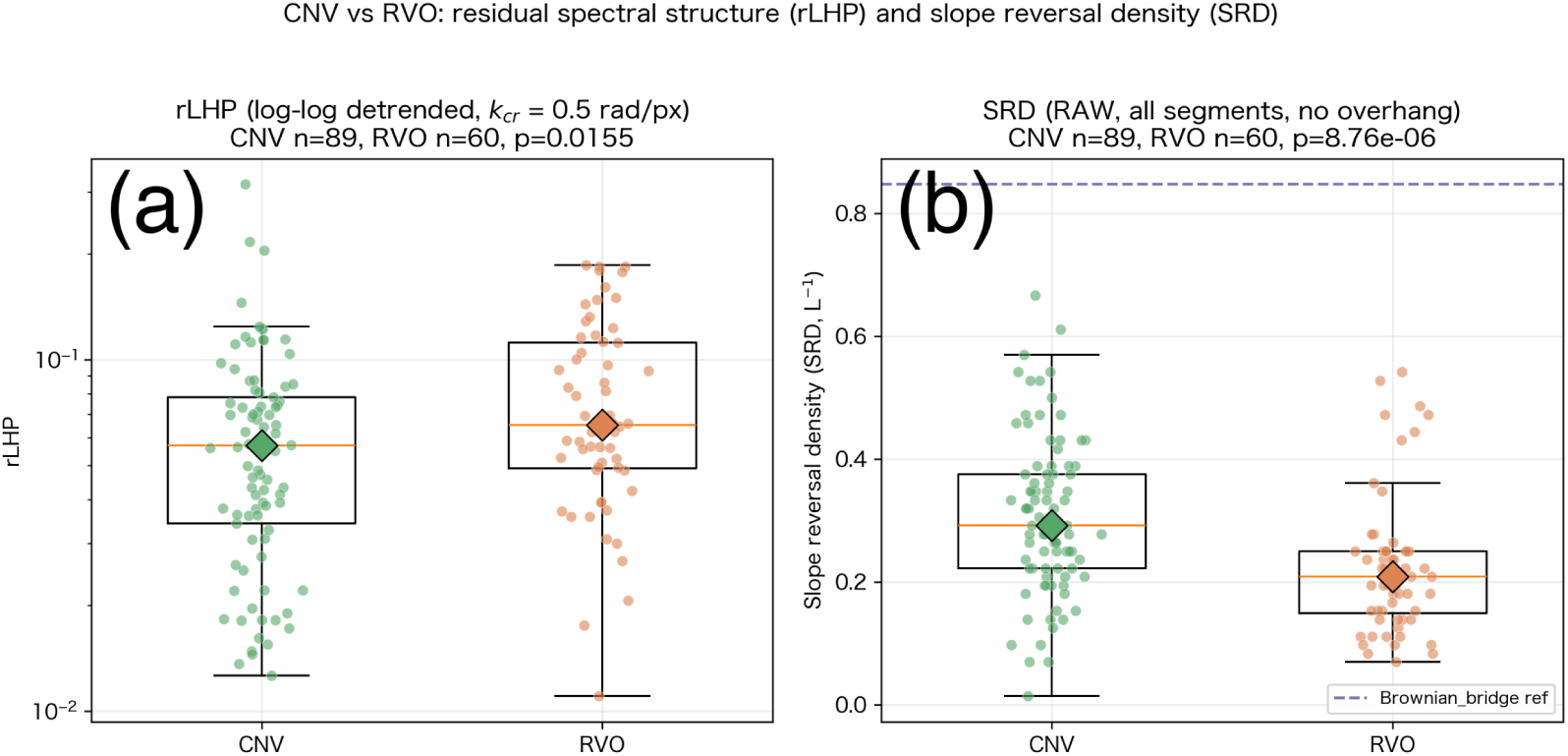
Segment-level comparison of CNV and RVO vessel morphology. **(a)** Residual low-to-high wavenumber power ratio (rLHP) at *k*_cr_ = 0.5 rad pixel^−1^ (log scale). **(b)** Slope reversal density (SRD; pixel^−1^); dashed line, Brownian-bridge reference. Box plots, median and interquartile range; circles, individual segments; diamonds, group medians. *n* = 89 (CNV) and *n* = 60 (RVO); two-sided Mann–Whitney *U* test, *p* = 0.015 (rLHP) and *p* = 8.8 × 10^−6^ (SRD).

## 5. Discussion

### 5.1. Research summary

In this study, we compared two mechanisms of vascular tortuosity—mechanically driven buckling (Euler–Bernoulli) and angiogenesis-driven curvature (Chap-lain–Anderson)—from a common spectral viewpoint. Buckling generates a characteristic peak in the power spectrum due to the selective growth of a dominant wavelength, consistent with observations in coronary vessels. In contrast, angiogenesis produces no fixed wavelength and exhibits an approximate (frequency)^−2^ scaling, consistent with tumor vasculature. We further proposed a two-phase scenario in which angiogenesis first generates irregular vessel paths, followed by mechanical remodeling that amplifies or regularizes curvature. Motivated by this framework, we introduced two quantitative metrics: the residual low-to-high wavenumber power ratio (rLHP), which quantifies a localized excess near a buckling-related wavenumber, and the slope reversal density (SRD), which measures the frequency of sign changes in local slope and is therefore higher for random-walk-like trajectories and lower for coherent buckling.

### 5.2. Other curved-structure formation mechanisms

Examples of meandering boundaries are known in other biological systems. For instance, cranial sutures form complex curved interfaces during skull growth; proposed models include stochastic interface growth (Eden front (Oota et al., 2004)), signal-factor-driven interface instability (Miura et al., 2009), and diffusion-limited aggregation (DLA)-based fractal growth (Zollikofer and Weissmann, 2011). These studies suggest that, beyond vessel-specific mechanisms, generic pattern-forming processes at growing interfaces can also generate tortuous geometries. Anatomical structures other than blood vessels that exhibit tortuosity include the vas deferens, cystic duct, and sweat glands(Ross and Pawlina, 2016). Their formation mechanisms remain poorly understood.

### 5.3. Possible application to clinical diagnostics

Retinal vascular tortuosity has been explored as a potential imaging biomarker for ocular and systemic diseases, such as cardiovascular diseases and stroke (Kalitzeos et al., 2013). For example, inflammatory diseases such as retinal vasculitis can be accompanied by increased vessel tortuosity (Zhang et al., 2025). Numerous automated methods for quantifying retinal-vessel tortuosity from fundus or OCTA images have been proposed (Ramos et al., 2019; Á et al., 2024). A common definition of tortuosity is the path-length-to-chord-length ratio, which is sensitive for distinguishing curved versus nearly straight segments; however, it is largely insensitive to *scale-dependent* signatures such as a dominant wavelength (buckling-like) or low-frequency power-law scaling (angiogenesis-like). The spectral profiles may provide complementary information for distinguishing *mechanistic causes* of vessel curvature from vascular images.

### 5.4. Effect of tortuous vessels

Tortuous blood vessels can have both physiological roles and pathological effects. Concerning the physiological role, anatomically, vessels around joints may form a curvature during axial shortening or follow inherently curved paths with excess length to prevent rupture during joint flexion and extension (Wensing et al., 1995). A tube, in general, may have additional roles. For example, the coiling of the epididymal duct has been proposed as a space-filling strategy that accommodates an extremely long tubular structure within the confined volume of the scrotum (Nelson, 2016). From an engineering point of view, curved tubes are often used to increase the performance of heat exchangers, like radiators (Naphon and Wongwises, 2006).

Tortuous vessels are also associated with several pathological consequences, including increased flow resistance, larger pressure drops, and reduced downstream perfusion(Wang et al., 2016). In addition, vessel curvature can induce disturbed hemodynamics, promoting endothelial injury and increasing the risk of atherosclerosis (Chiu and Chien, 2011) and the formation of blood co-agulants (Sunderland et al., 2022). Tortuosity can also complicate catheter-based interventions(Konigstein et al., 2021).

### 5.5. Future directions

At present, our original goal of inferring etiology from the morphological features of tortuous retinal blood vessels has not yet been fully achieved. In future work, we will improve the image processing method to detect the differences in pathogenesis.

Currently, we confine the theoretical framework to two models: the Chaplain-Anderson model and buckling; however, other mechanisms may also be possible. For example, it is known that nail bed capillaries show a twisted structure in scleroderma patients (Smith et al., 2023). In this case, the capillary becomes unstable, so a different type of model may be needed. In the case of cerebral vessels, normal vessels show a twisted structure, probably due to gyrification (Kulenović and Dilberović, 2004; Padget, 1948).

## Acknowledgements

This work was supported by the Japan Society for the Promotion of Science (JSPS) KAKENHI Grant Number 24K02036.

## CRediT authorship contribution statement

**Hiroto Kawanaka:**Conceptualization, Methodology, Software, Validation, Formal analysis, Investigation, Data curation, Visualization, Writing – original draft. **Takashi Miura:** Conceptualization, Methodology, Supervision, Project administration, Funding acquisition, Writing – review & editing.

## Declaration of competing interest

The authors declare that they have no known competing financial interests or personal relationships that could have appeared to influence the work reported in this paper.

## Data availability

The OCTA500 retinal vessel dataset used in this study is publicly available at https://ieee-dataport.org/open-access/octa-500 (Li et al., 2024). Analysis code and processed summary data supporting the findings are available from the corresponding author upon reasonable request.

## Notes

### Competing Interest Statement

The authors have declared no competing interest.

### Summary of Updates

Added quantitative measures to distinguish pathology.

